# Accurate Identification of Extrachromosomal Circular DNA from Long-read Sequences

**DOI:** 10.1101/2022.05.13.491700

**Authors:** Visanu Wanchai, Piroon Jenjaroenpun, Thongpan Leangapichart, Gerard Arrey, Charles M Burnham, Maria C Tümmle, Jesus Delgado-Calle, Birgitte Regenberg, Intawat Nookaew

## Abstract

Extrachromosomal circular DNA (eccDNA) of chromosomal origin is found in a range of eukaryotic species and cell type including cancer where eccDNA with oncogenes appear to drive tumorigenesis. Most studies of eccDNA employ short-read sequencing to identify for their identification. However, short-read sequencing cannot resolve the complexity of genomic repeats, which can lead to missing eccDNA identification. An alternative is the long-read sequencing technologies that can potentially be used to construct complete eccDNA. We present a software suite, **<u>C</u>**onstruction-based **<u>R</u>**olling-circle amplification for eccDNA Sequence **<u>I</u>**dentification and **<u>L</u>**ocation (CReSIL) 2.0, to identify and characterize eccDNA from long-read sequences. CReSIL’s performance in the identification of eccDNA, with a minimum F1 score of 0.98, is superior to the other bioinformatic tools based on simulated data. CReSIL provides many useful features for genomic annotation, which can used to infer eccDNA function and Circos visualization for eccDNA architecture investigation. We demonstrated CReSIL’s capability in many of the long-read sequencing datasets. This includes datasets enriched for eccDNA as well as whole genome datasets from many cells that contained large eccDNA. CReSIL suite software will be a versatile tool to deeply investigate eccDNA in biological samples.

## INTRODUCTION

Extrachromosomal circular DNA (eccDNA) of chromosomal origin is a common across the eukaryotic kingdoms from plants to fungi and animals^1-18^. EccDNA can range in size from a hundred base pairs to several megabases and thereby often capture genes or parts of genes^5, 19-21^. A lack of centromeres on eccDNA means that these elements segregate unfaithfully and have a tendency to accumulate in subpopulations of cells that can thrive on a high copy number when genes on the eccDNA is expressed and provides a selective advantage^20^. This is particularly evident in the unicellular yeast cells and cancer cells. Yeast cell grown under nutrient limiting conditions can obtain a selective advantage when transporter genes are trapped and accumulate in clonal sub-populations^5, 22^. An example of this is observed when yeast cells are grown with limited glucose and clones of yeast cells with circularized glucose transporter genes [*HXT6/7*^*circle*^] on eccDNA accumulate in the population^5^. Tumor cells accumulate oncogenes on circular DNA that is often megabase sized (ecDNA)^20, 23^. These structures are often complex and comprised of several fragments, such as enhancers and several oncogenes, suggesting that ecDNA can also evolve to become more transcriptionally efficient ^24, 25^. These recent advances in our understanding of eccDNA are mainly obtained through short read sequencing with tools such as AmpliconArchitect^26^, Circle-Map^27^, ECCsplorer^28^ and ccDNA_finder^29^.

However, identification of eccDNA from short reads are limited by the capacity to identify repetitive regions of genomes such as centromeres and complex eccDNA composted of several fragments. This problem can be overcome with long read sequencing technologies such as Pacific Bioscience Technology (PacBio) and Oxford Nanopore Technology (ONT) because the long reads span longer stretches and thereby spanning elements in repeats and multiple fragments in complex eccDNA. Identification is further aided by amplification of the eccDNA with Rolling Circle Amplification (RCA)^30-32^. This simple yet powerful approach allows for reads that have often traveled around the circle several times, generating concatemeric tandem copies (CTC). Still bioinformatics tools to identify eccDNA from RCA long read sequences are limited and the tolls have never been compared. Besides, Metha et. al. presented CIDER-Seq2 that is designed for characterization of virus genomes^33,34^.Zhang et al presented ecc_finder that is made for both short and long read sequences^29^. The tool was extensively tested in short-read sequences; however, the performance evaluation on long-read dataset is limited. Wang et. al. presented the eccDNA_RCA_Nanopore software^15^. This tool focusses only on sequences contain CTC and ignored the remaining 52% of the sequenced reads which could be the important products derived from incomplete RCA of large size eccDNA or DNA breakages during the experimental procedures. We have recently presented two tools for mapping of eccDNA, NanoCircle and CReCIL^16^. NanoCircle identifies complex and simple circles from sequence reads that span junction site in the circular DNA, but often fail to correctly assemble complex circles. CReSIL designed to construct eccDNA de novo, however advancing computational workflow and additional features aid deep investigations of the identified eccDNAs will need to be developed.

Here we present an improved bioinformatics software suite, <u>C</u>onstruction-based <u>R</u>olling-circle amplification for <u>e</u>ccDNA <u>S</u>equence <u>I</u>dentification and <u>L</u>ocation 2.0 (CReCIL 2.0) to accurately construct and represent eccDNA from long read sequences. We benchmarked it to other known tools for identification of eccDNA from long read sequences and find that it is superior to all other available tools. CReSIL provides useful features to investigate genomic annotations, sequence variations of eccDNA as well as visualization of eccDNA architecture. We demonstrated the capability of CReSIL to identify eccDNA molecules from many human and mouse samples enriched for eccDNA and we finally showed how CReCIL can be used to identify eccDNA from whole genome long-read sequencing (WGLS) datasets.

## RESULTS

### EccDNA enrichment workflow and CReSIL computational workflow for eccDNA identification

We initially designed a computational workflow for identification of eccDNA from long read sequence samples enriched for eccDNA (Fig. 1.0). Enrichment was obtained by removal of chromosomal DNA and mitochondrial DNA from eccDNA. In this procedure the mitochondrial DNA was linearized using either rare cutting restriction enzymes DNA or CRISPR-Cas9 specifically designed to target mtDNA; after which the linear chromosomal and mitochondrial DNA was digested using Exonuclease V. We next amplified the eccDNA molecules using RCA, which allowed for DNA sequences that spanned the eccDNAs more than once. The RCA approach produced hyperbranched DNA products that required de-branched with the T7 endonuclease before sequencing library preparation to obtain long reads and minimize pore clogging during DNA sequencing that would otherwise result in ghost sequences (i.e., improperly generated sequences that are unmappable or have similarities to the known sequence but in the opposite orientation). ONT sequencing was next performed to generate long-read sequences that were used as the input for the CReSIL 2.0 bioinformatic analysis.

**Figure 1.**
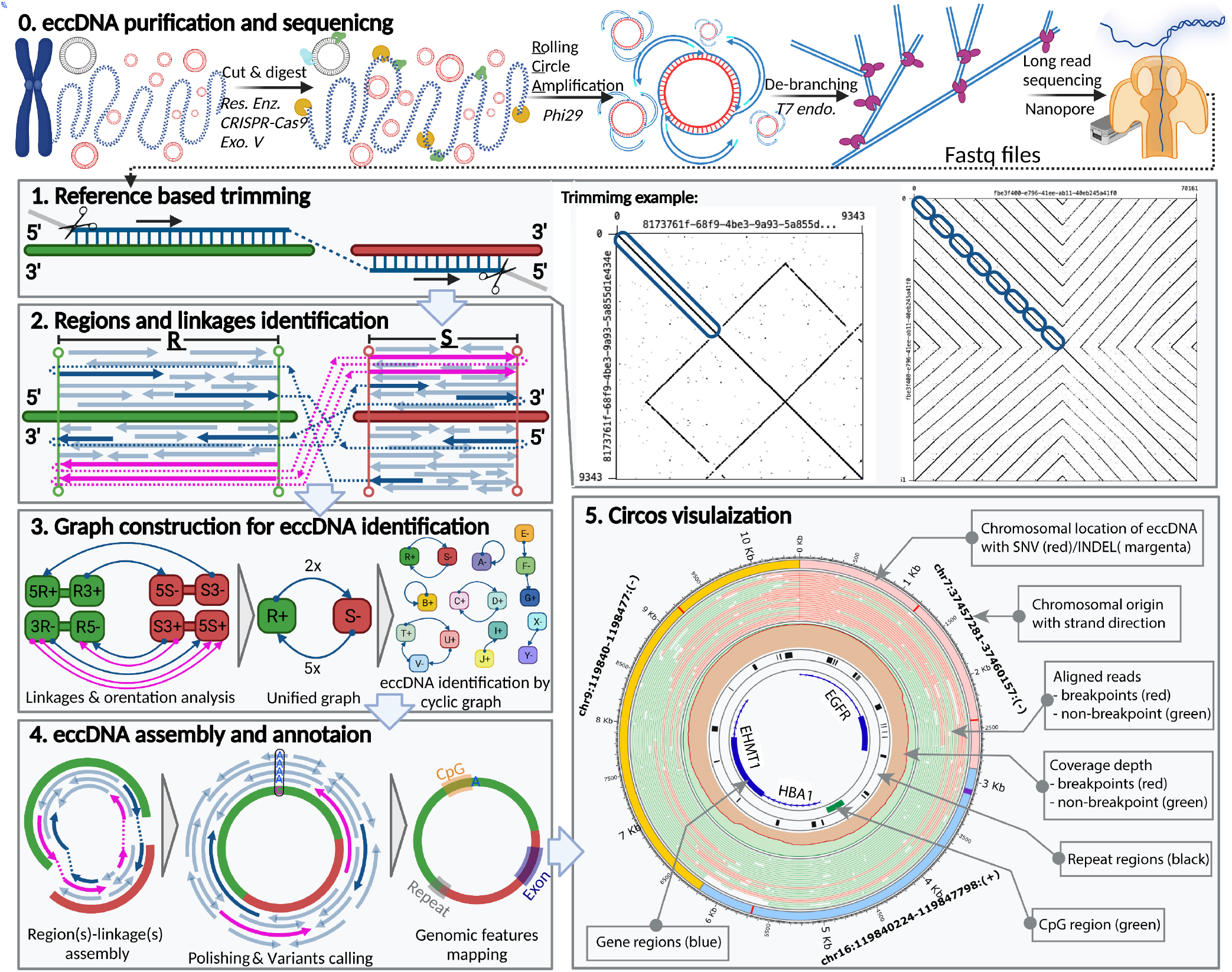
EccDNA identification by long-read sequencing. **Step 0**. Experimental workflow beginning with purified genomic DNA; chromosomal DNA (blue), mitochondrial DNA (magenta), eccDNA (red), restriction enzyme (green), CRSPR-Cas9 (cyan), exonuclease V (yellow). **Step 1**. A read (left panel) can be aligned on regions of 2 chromosomes (green and red) of the reference genome (blue) with breakpoint reads (dashed line) linking the aligned chromosomal regions and trimmed unmapped portions (gray). Self-read dot plots (right panels) showing read regions that align to the reference genome (blue ovals); reads without CTCs (left dot plot) and with CTCs (right dot plot). **Step 2**. Merged regions R and S with the reference sequences (green and red bars) and the breakpoint event (dashed lines) that links the two regions; arrows for non-breakpoint reads (light blue), breakpoint reads (blue), and reads with CTCs (magenta) point in the direction of the aligned orientation on the plus strand of the reference sequence (arrows above the bars) or the minus strand (arrows below). **Step 3**. The information of the read alignments were converted to directed graphs. **Step 4**. The reads were assembled using our developed regions/linkages algorithm; the assembled sequences were polished and variants were identified; and eccDNA was annotated with genomic features such as exon, repeat and CpG. **Step 5**. Circos visualization to present the eccDNA architecture.

The long-read sequences generated from the procedure above can be classified into three categories, which are break point read without CTC, CTC read and normal read. Break point read without CTC (blue arrows in Fig. 1.1) is the read contains no CTC that can be aligned on reference sequence more than one region. This provides important evidence to link regions together (blue dashed lines in Fig. 1.1) for circle construction. CTC read (magenta arrows in Fig. 1.1) is the read contains CTC derived from RCA amplification product that spanned the eccDNAs more than once. Therefore, CTC read always contains break point(s) (magenta dashed lines in Fig. 1.1). The normal read (light blue arrows in Fig. 1.1) is the read can be aligned only one region on reference sequence, therefore, it contains no breakpoint.

Even though the de-branching was performed, we observed ghost sequences in the long-reads data as examples of dot plots illustrated in Fig. 1.1. Therefore, Step 1 of CReSIL was the preprocessing of the raw reads by reference-based trimming and is essential in avoiding false positives from remaining ghost reads. First, the reads were aligned on the reference genome. Next, high confidence mapped region(s) of the reads were obtained and the rest were trimmed out. For example, the read without CTC (left dot plot of Fig. 1.1) has a sequence length of 9,343 nucleotides (nt); we found a 3,514 nt region at the beginning of the read that was mapped to the reference genome. CReSIL trimmed the rest of the read, which was assumed to be a ghost sequence of self-inverted repeats followed by an unmapped sequence. An example of a CTC read (right dot plot of Fig.1.1), was a sequence of 70,161 nt. We found that the first half of the read contained 8 consecutive sequence copies that could be mapped on the same region of the reference genome. The second half of the read was a sequence of a self-inverted repeat, which was assumed to be a ghost sequence, that was discarded by CReSIL. The reference alignment results of the individual reads were used for the next step.

In Step 2 (Fig. 1.2), chromosomal aligned locations of the trimmed reads were aggregated to identify merged regions representative of the chromosomal origins of eccDNA. On the individual merge regions, we typically observed 3 types of aligned reads: 1) reads aligned only on 1 region (normal reads), 2) reads aligned on multiple regions without CTC (break point reads without CTC), and 3) reads aligned on multiple regions with CTC (CTC reads). CReSIL recorded the aligned reads of the identified individual merged region, the linkages of the identified merged region(s), the orientation of the reads (indicated by arrows direction), and the aligned strand orientation of the chromosomal regions. The last two types contain breakpoint(s), which is the important evidence to identify linkages that connect region(s) together.

In Step 3 (Fig. 1.3), CReSIL formulated the recorded regions and linkages information into graph representations. First, CReSIL constructed directed graphs with the information of regions, terminals, and strands; therefore, an individual region contained 4 nodes and multiple edges derived from linkages (Fig. 1.3. left panel). After the linkages and orientation analysis based on read alignment orientation and strand, the graphs were unified, resulting in one node representing one region and the weight of the edges representing the number of linkages (Fig. 1.3. middle panel). From this point, CReSIL identified high-confidence eccDNA from cyclic graphs and low-confidence eccDNA from acyclic graphs.

In Step 4 (Fig. 1.4), CReSIL performed region(s) and linkage(s) assembly based on the graphs (Fig. 1.4. left panel). All the reads belonging to an individual graph were assembled and were the polished to generate a consensus sequence. During polishing, CReSIL also identified mutations present in the eccDNA (Fig. 1.4. middle panel). CReSIL annotated some selected genomic features on the identified eccDNA (e.g., exon, intron, repeat, CpG).

In Step 5 (Fig. 1.5), CReSIL generated an input file of selected eccDNA for visualization to illustrate eccDNA architecture using Circos software^35^. A plot contains information about the chromosomal origin of eccDNA; any single nucleotide variation (SNV) or an insertion-deletion mutation (INDEL); a read alignment; and coverage of breakpoint reads, non-breakpoint reads, and selected genomic features.

### CReSIL outperformed in eccDNA detection compared with other tools

To evaluate the performance of CReCIL, we generated a synthetic eccDNA datasets of 1,300 true positives, which mimic long-reads derived from eccDNA purification and RCA amplification (Fig. 1.0), and 1,300 true negatives by randomly selecting chromosomal regions of the human genome (hg19) to evaluate the performance of CReSIL in comparison with the other tools such as eccDNA_RCA_nanopore^15^, ecc_finder^39^, and the long-read de novo assembler Flye^36^. We simulated eccDNA following the size and region distribution previously reported by Henriksen et al^16^ (Fig. 2A). The size distribution of the simulated eccDNA true positive and true negative datasets were similar (Fig. 2A) with a minimum size of approximately 500 nt, a maximum size of approximately 34,000 nt, and a median size of approximately 6,200 nt. The simulated sets contained both simple and complex (> 1 region) eccDNA with a similar distribution of chromosomal-containing region (Fig. 2B), and varying depth coverage from 3x to 100x. We used PBSIM2^37^ software to simulate long-reads based on the simulated eccDNA sets varying depth coverage from 3x to 100x. Considering the true positive dataset, we found that in the lower sequencing depth, the number of simulated eccDNA that contain CTC reads was reduced (Fig. 2C) due to lower probability to generated CTC reads by PBSIM2 that mimicked the incomplete circular amplification and/or DNA breakage events.

**Figure 2.**
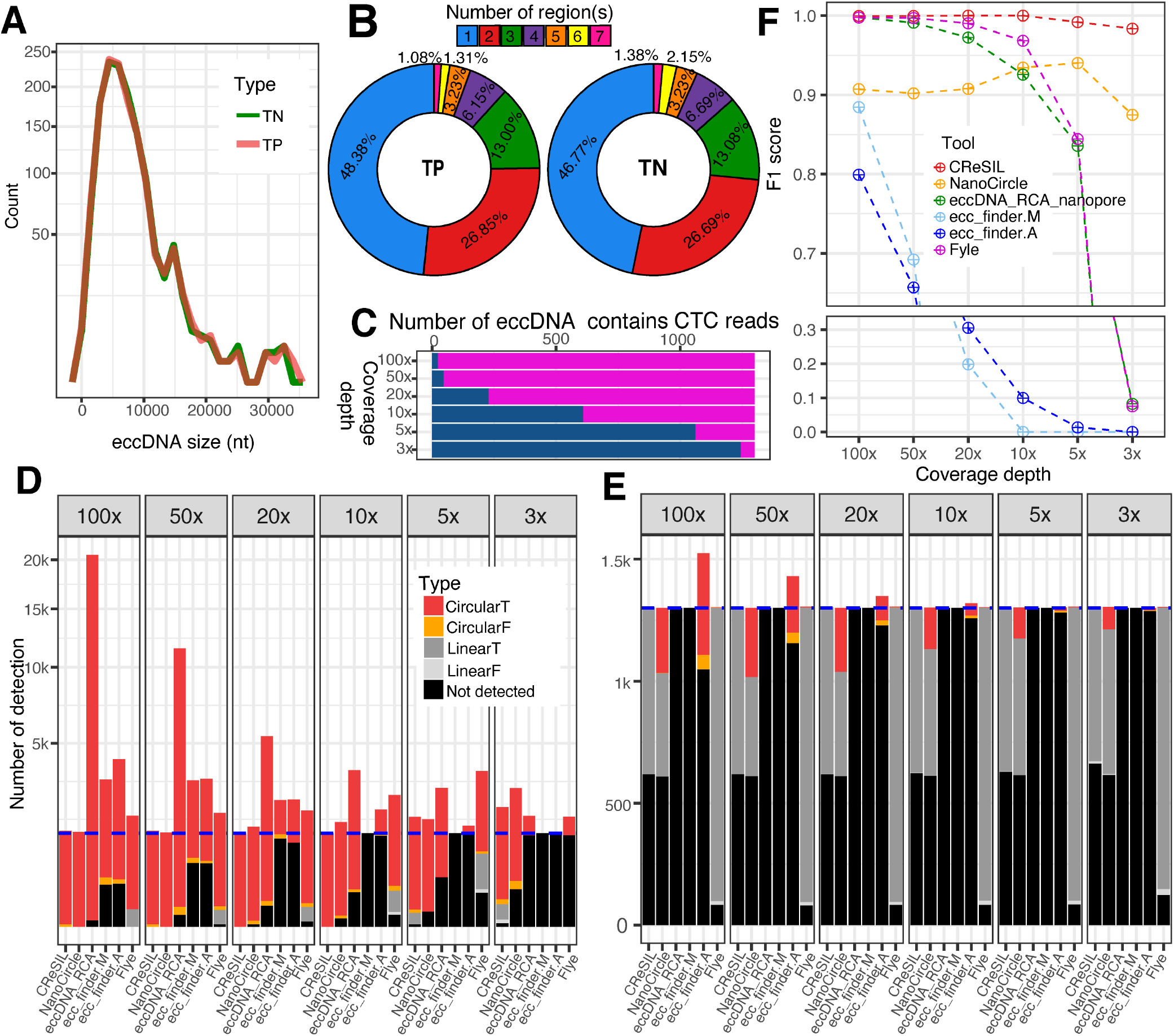
Evaluation of eccDNA detection performance of CReSIL and its comparison with the other tools. **A**) Frequency polygon plot showing the size distribution of simulated eccDNA of true positive (TP, orange line) and true negative (TN, green line) datasets. **B**) Donut plots showing the percent distribution of the number of chromosomal regions of the individual simulated eccDNA datasets. **C**) Bar plots showing the number of simulated eccDNA that contains CTC reads (magenta) and non-CTC reads (dark blue) across different sequencing depths. **D**) Stacked bar plots showing the number of eccDNA detected for the true positive dataset by different tools. The eccDNA detection results are classified into 5 categories; CircularT (red) = circular sequences with 95% reciprocal overlaps and 90% identity with the simulated eccDNA sequences, CircularF (orange) = circular sequences without the criteria, LinearT (dark gray) = linear sequences with the criteria, LinearF (light gray) = linear sequences without the criteria, and not detected (black) = the tool cannot detect; number of true positive eccDNA (blue dashed line). **E**) Stacked bar plots presenting the number of eccDNA detected for the true negative dataset. See the color code for panel D, number of true negative eccDNA (blue dashed line). **F**) Point and line plots showing the performance (F1 score) of eccDNA detection of the individual tools across different sequencing depth.

The detection of eccDNA results obtained from CReSIL and the other tools based on true positive (Fig. 2D) and true negative (Fig. 2E) datasets. CReSIL and NanoCircle, correctly detected almost every true positive datasets except low coverage of 5x and 3x (Fig. 2D). At 100x, eccDNA_RCA_nanopore detected a large amount of eccDNAs of approximately 20,000 eccDNAs, which is high redundancy derived from individual CTC reads. Because eccDNA_RCA_nanopore is designed to detect eccDNA derived from CTC reads only; eccDNA_RCA_nanopore missed detection of eccDNA whose simulated reads contained no CTC reads. We also compared to ecc_finder in 2 modes, assembly(ecc_finder.A), and mapping(ecc_finder.M). The eccDNA detection results derived from the 2 modes revealed a high fraction of true positives that were not detected at high depth coverage of 100x and 50x and missed detection of almost every eccDNA at low sequencing depth of 10x, 5x and 3x. The de novo assembly tool, Flye, performed well in the detection of eccDNA at 100x (missed detection of 48 eccDNAs), yet the performance of Flye dropped when the sequencing depth was reduced. Considering the true negative datasets (Fig. 2E), almost every tool controlled the false positive rate very well, except ecc_finder assembly mode(ecc_finder.A), which produced high number of false positives at 100x and reduced as the sequencing depth decreased. NanoCircle has similar number of false positive across all sequencing depth, which derived from complex circles, only. We next calculated the F1 score to compare the eccDNA detection performance across the tools at the various coverage depths (Fig. 2F, the precision and recall of individual tool and scenario is provided in Supplementary Table S1.). All tools performed very well at 100x, but F1 scores dramatically dropped when sequencing depth was reduced, except CReSIL, which maintained an F1 score of 0.98 at 3x. This indicated the outperformance of CReSIL in eccDNA detection from long-reads derived from eccDNA enrichment and RCA amplification over other available tools.

### Identification and characterization of eccDNA from eccDNA enrichment samples using CReSIL

To demonstrate the capabilities of CReSIL on real data, we next performed eccDNA enrichment and long-read sequencing following the experimental workflow as shown in Fig. 1.0 to generate high quality datasets of 3 human multiple myeloma cancer cell lines, APR1, EJM, and JJN3; 2 mouse tissues, pancreas and cortex; and 3 mouse cells, MLOY4 (osteocyte-like), 5TMG1 (multiple myeloma), and E0771 (breast cancer). In addition, we included published eccDNA enrichment long-read sequencing data of a human sperm dataset from Henriksen et al^16^ and the mouse embryonic stem cell mESC dataset from Wang et al^15^ in the analysis

The datasets ranged from 0.25 to 4.16 million reads after trimming (Fig. 3A), CResIL detected a variety of reads containing CTC with debranching ranging from 9.4% to 53.7% and only1.8% were CTC in samples that had not been debranched prior to sequencing. This indicated the importance of T7 endonuclease treatment before sequencing. After trimming, CReSIL keep high fraction of reads for further step with an average of 76% of reads for the CTC reads and 98% of reads for the normal reads (Fig. 3B).

**Figure 3.**
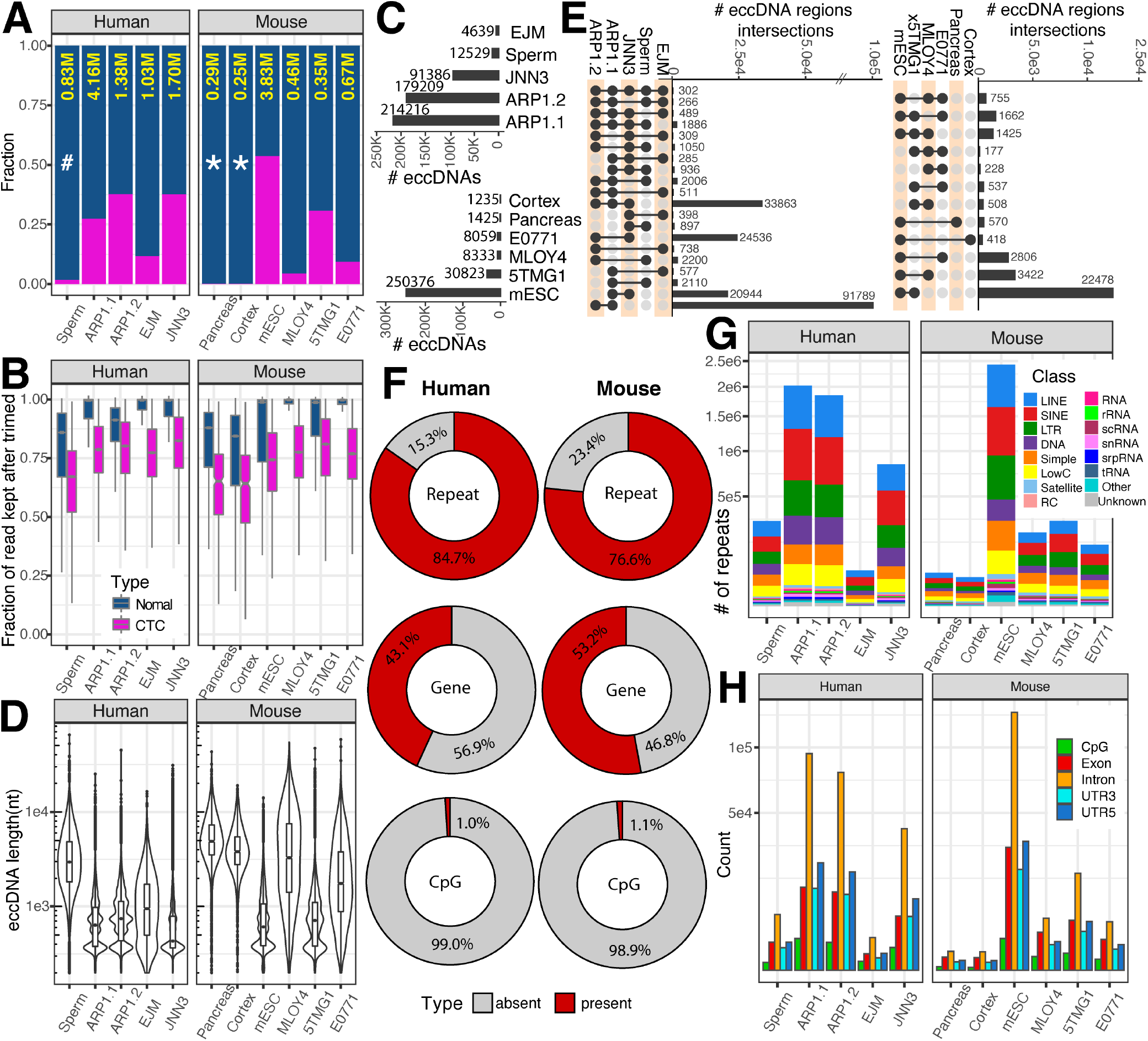
Identification of eccDNA from long-read sequencing of eccDNA enrichment samples derived from human and mouse cells. **A**) Stacked bar plot showing the fraction of non-CTC reads (dark blue) and CTCs (magenta) across 5 human cell samples and 6 mouse cell samples; total number of high-quality reads after the CReSIL trimming step (yellow), in million read units; * = datasets prepared by primer-free based RCA; the rest were prepared by primer-based RCA; # = sample prepared for sequencing without debranching. **B**) Box-whiskers plots showing the fraction of reads used for eccDNA identification that were kept after trimming of normal (dark blue) and CTC (magenta) reads. **C**) Bar plots summarizing the number of eccDNA in human (top panel) and mouse (bottom panel) datasets **D**) Violin-boxplots showing the distributions of length of identified eccDNA. E) Upset plots showing overlaps (right panels), only overlapped numbers over 100 is shown. **F**) Donut charts showing the percentages of eccDNA harbored repeats, genes, and CpG islands. **G**) Stacked bar plots showing the frequency of different classes of repeats harbored in identified eccDNA. **H**) Bar plots showing the frequency of CpG, exon, intron, 3’UTR, and 5’UTR harbored in identified eccDNA.

In human datasets (Fig. 3C top panel), CReSIL identified a high number of eccDNA in the cancer cell lines, over 200,000 eccDNAs for both replicates of APR1 and over 90,000 eccDNA for JJN3 cells. CReSIL also identified over 12,000 eccDNA for sperm and over 4,600 eccDNA for EJM cells. In mouse datasets (Fig. 3D bottom panel), CReSIL identified over 250,000 eccDNAs from mESC, over 30,000 for 5TMG1, over 8,000 eccDNAs for MLOY4 and E0771 cells, and less than 1,500 eccDNAs for the 2 tissues. Overall, of the human and mouse dataset, most of the identified eccDNA were formed from a single region in the genome (simple eccDNA) and only an average of 1.5% were complex eccDNA made of several fragments of chromosomal DNA (see example Fig. 1 step 5 and Supplementary Table S2). The length size distribution of the identified eccDNA of individual datasets is shown Fig. 3D. The smallest size of eccDNA is 200 nt, which is the same as the minimum read length of input to CReSIL. We recorded the biggest size of eccDNA to approximately 65,000 nt in sperm for the human dataset and approximately 58,000 nt in E0771 for the mouse dataset. Next, we checked the overlaps among the identified eccDNA for each organism as shown in the upset plots (Fig. 3E). We observed relatively low numbers of overlaps comparing the numbers of identified eccDNA. This reflected a randomness of eccDNA biogenesis as proposed in Møller et. al.^8, 12^ and Wang et al^15^. However, we found over 90,000 recurrent eccDNA between replicates of the ARP1 cell line (∼50%), indicating a cell specific population of eccDNA that possibly benefits the ARP1 cell line (Fig. 3C top right panel).

Characterization of genes and genomic contents harbored by eccDNA is essential for understanding the phenotypic effect a specific eccDNA might have in tumor cells^24, 38^ or other cells where eccDNA can provide a selective advantage^5, 22^. Therefore, CReSIL also provides a function for genomic annotations of individual eccDNA. We annotated three types of genetic elements, repeats, genes, and CpG islands, on the identified eccDNA. CReSIL mapped the repeats on a high percentage of identified eccDNA of 84.7% in the human datasets and 76.6% in the mouse datasets (Fig. 3E), which is similar to the previous reports using short-read sequences to identify eccDNA^12, 17^. For genes, 43.1% of the identified eccDNA in the human dataset and 53.2% in the mouse dataset harbored genes or parts of genes (Fig. 3E middle panel). We observed very small amount, less than 1.5% of the identified eccDNA harbored CpG islands (Fig. 3E bottom panel). Repetitive sequences were commonly found in eccDNA reported in many studies^12, 17, 18^ and could promote circles forming through microhomology^18, 39, 40^. The 16 different repeat classes were distributed across individual datasets (Fig. 3F). Long interspersed nuclear elements, short interspersed nuclear elements, and long terminal repeats were the most frequently found classes in the identified eccDNA. Of the repeat types, the introns were most frequently found in identified eccDNA harbored genes or part of genes. We observed the high frequency of exons were harbored by the identified eccDNA (Fig. 3G).

Because CReSIL identifies eccDNA by aggregation of all reads in a sample and calculates the coverage depth of individually identified eccDNA, we can use this information for qualitatively ranking the relative abundance of eccDNA molecules. Based on the hexbin plots (Fig. 4A), we did not observe a correlation between the size of the identified eccDNA and their coverage depth for both human and mouse datasets. Interestingly, we found that the most frequent coverage depth of the identified eccDNA is approximately 10x for both datasets. Next, CReSIL generated consensus sequences of identified eccDNA and called genetic variations for individually identified eccDNA. At the variant quality score cut off of 20, we found an average of 35.6% of the identified eccDNA for the human dataset and 32.0% for the mouse dataset have variant(s) (Fig. 4B). Single nucleotides variation event gave the highest fraction of the variants, except for mouse pancreas and cortex eccDNA, in which deletion event gave the highest fraction of the variants (Fig. 4C). We observed a low fraction of insertion events across all datasets.

**Figure 4.**
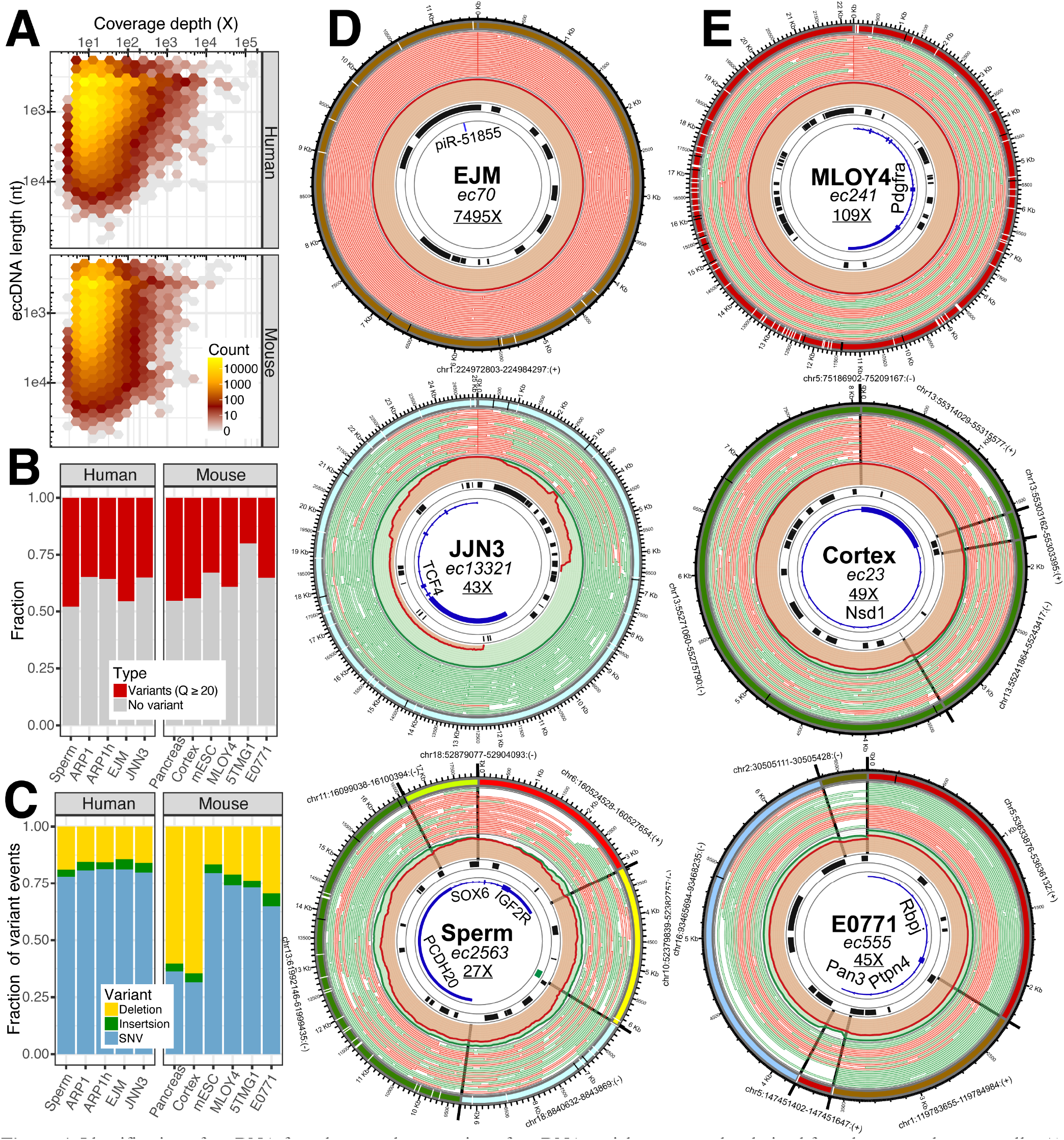
Identification of eccDNA from long-read sequencing of eccDNA enrichment samples derived from human and mouse cells. **A**) Hexagonal bin plots showing no correlation between the length of the identified eccDNA with their coverage depth. **B**) Bar plots showing the fraction of the identified eccDNA containing variants (SNVs and/or INDELs) based on a quality cutoff of 20. **C**) Bar plots showing the distribution of deletions, insertions, and SNVs of the identified eccDNA. **D**) Circos plots visualizing the architecture of 3 selected human eccDNA datasets. **E**) Circos plots visualizing the architecture of 3 selected mouse eccDNA datasets (see Fig. 1.5 for lane annotation; data set information (bold), eccDNA name (italics), coverage depth (underline).

CReSIL also provides functions to make Circos plots visualize the architecture of identified eccDNA. We arbitrarily selected 6 showcases of eccDNA from the human (Fig. 4D) and mouse datasets (Fig. 4E). Of the human showcases, identified from the EJM dataset, ec70 has a size of approximately 11,500 nt and has an extremely high coverage depth of almost 7,500x, indicating a very high abundance of this eccDNA molecules. Ec70 harbored only a short non-coding RNA, piR-51855. Most of the reads contributing to the identification of ec70 were CTC reads. In contrast, identified from the JJN3 dataset, ec13321 has a big size of approximately 25,000 nt which is based on non-CTC reads with strong support of breakpoint reads. Ec13321 had 43x coverage depth and harbored 5 exons of the transcription factor 4 gene. Lastly, identified from the sperm dataset, ec2563 is a complex eccDNA that comprises 6 regions of different chromosomes with a total length of approximately 18,000 nt and harbors a CpG island and the parts of 3 genes, insulin growth factor 2 receptor, sex-determining region Y–box transcription factor 6, and protocadherin 20, which came from different chromosomal origins. Of the mouse showcases, ec241 was identified from the MLOY4 dataset and is approximately 22,000 nt with a high coverage depth of 109x. Interestingly, ec241 contains 5 exons of the platelet-derived growth factor receptor alpha gene, which is a well-known marker for mesenchymal lineage cells^41^, reflecting the origin of the osteocyte-like MLOY4 cells. Identified from the cortex dataset, ec23 is a complex eccDNA with a length of approximately 8,000 nt and 4 regions from chromosome 13. Interestingly, based on the chromosomal coordinates, ec23 began at chr13:55303162 and ended at chr13:55315577 (12,416 nt). Ec23 showed that 4 regions within the chromosomal coordinates were combined to generate an eccDNA with a size of 8,068 nt, which is a substantial deletion (a total of 4,348 nt). Ec23 had 49x coverage depth and harbored 1 exon of the nuclear receptor binding SET domain protein 1 gene, which is known to be associated with Sotos syndrome^42^ (caused by mutations in this gene). Finally, identified from E0771, ec555 is a complex eccDNA of 5 regions from 4 different chromosomes and has size of approximately 6,600 nt with 45x coverage depth. Ec555 harbored the parts of 3 genes, recombination signal binding protein for immunoglobulin kappa J region, protein tyrosine phosphatase non-receptor type 4, and poly(A) specific ribonuclease subunit. The evidence used to construct the molecule derived from both CTC and normal reads with a similar fraction of half.

### Identification of eccDNA from long-read whole genome sequencing datasets by CReSIL

Turner et al reported that approximately 50% of the focal copy number amplification in tumors derive from eccDNA molecules^20^. The structure can be constructed by their sequence architecture and by the AmpliconArchitect algorithm and validated some cases by long-read sequences, reported by Deshpande^26^. Therefore, we extended CReSIL capability to identify eccDNA molecules from WGLS by implementing an additional workflow (Fig. 5A) to identify focal regions and their linkages then fed it to the graph construction algorithm of CReSIL. The reads were trimmed by CReSIL. Beginning with the read alignment result, CReSIL calculated the coverage depth of small windows of 10 nt across the reference chromosomes to ensure that the signal of a small-sized eccDNA was captured. The average coverage depth of individual chromosomes was recorded to estimate the backgrounds, which was used to subtract the coverage depth of all windows with respect to their chromosomal location. In parallel, CReSIL identified the breakpoint positions from the read alignment result. After the subtraction, the focal amplified regions were uncovered by aggregation of the windows that had depth higher than the background. The focal amplified regions were refined by the breakpoint locations. The refined regions and the linkages information were converted in the graph construction (Fig. 1.3) of the CReSIL workflow. Then the eccDNAs were identified through the CReSIL workflow.

**Figure 5.**
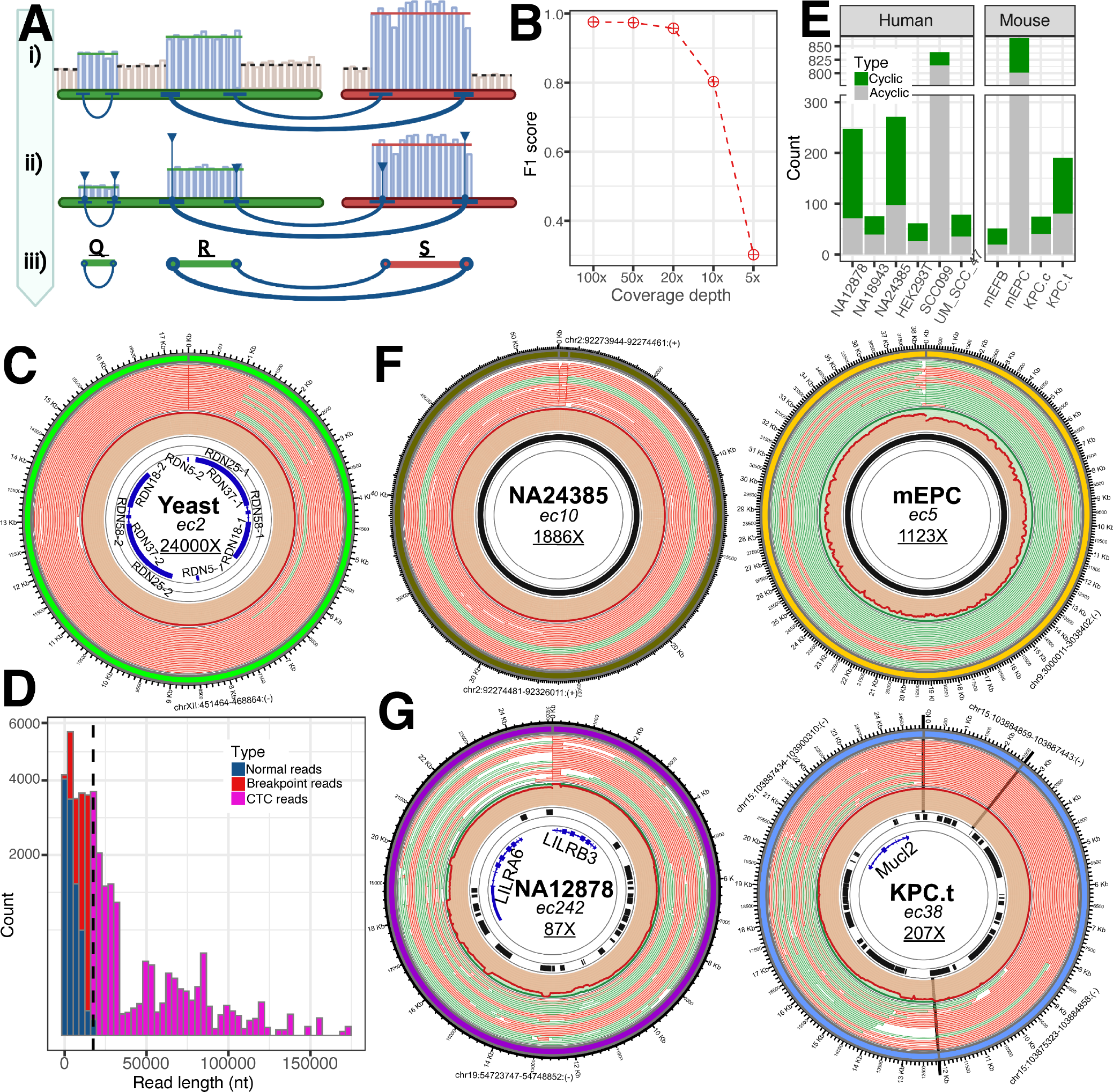
Identification of eccDNA from WGLS of yeast human and mouse cells. **A**) Scheme showing CReSIL extension workflow to identify eccDNA from WGLS dataset; the sequencing depth on a small window of focal amplified regions (blue bars) was higher than background (gray) and the identified focal regions (horizontal lines) are shown with the breakpoint reads generating linkages between the regions (dark blue curved lines); also shown are the number of breakpoints reads (thicker lines = more reads) and the number breakpoints reads at that location (higher vertical lines = more reads). **B**) Dot-line plot of F1 scores showing the performance of CReSIL in the identification of eccDNA from WGLS synthetic data by mixing 20x of human genome reads with different coverage of simulated eccDNA reads of true positive and true negative set. **C**) Circos plot showing the identified yeast eccDNA of the known circular rDNA. **D**) Histrogram showing the distribution of normal reads (darkblue), breakpoints reads (red), and CTC reads (magenta) with the size of identified eccDNA of rDNA (vertical dashed line). **F**) Circos plots showing examples of identified eccDNA with high coverage depth on centromere regions of human (left) and mouse (right) datasets. All are satellite repeats. **G**) Circos plots showing examples of identified eccDNA containing gene(s) of human (left) and mouse (right) datasets (see Fig. 1.5 for lane annotation; dataset information (bold), the eccDNA name (italic) and coverage depth (underline).

We assessed the capability of CReSIL to identify eccDNA from WGLS using this workflow. To mimic the situation, we used PBSIM2 to randomly generate long-read datasets of the human genome at 20x coverage depth and mixed them with the simulated eccDNA true positive and true negative datasets of the varying coverage depths. CReSIL has a very good performance, with an F1 score > 0.98 (Fig. 5B) when the coverage depth of simulated eccDNA was 100x, 50x, and 20x. The F1 score dropped approximately to 0.8 at 10x, and the performance was low (score of ∼0.3) at 5x.

We demonstrated the CReSIL workflow on the generated WGLS of *Saccharomyces cerevisiae* dataset of 4.3 Gb, corresponding to 356x genome coverage depth. CReSIL identified only 2 eccDNA with a very high coverage depth. Interestingly, one of them is the well-known circular ribosomal DNA (rDNA) molecule in yeast. CReSIL identified the eccDNA with a size of approximately 17,000 nt, covering the whole RDN gene clusters of RDN37, RDN25, RDN18, RDN58, and RDN5 (2 clusters based on the reference genome), with an extremely high coverage depth of 24,000x (Fig. 5C). The result agreed with the recent studies that reported the massive accumulation (> 99% in aged cells) of circular rDNA in yeast based on the circularization for in vitro reporting of cleavage effects by sequencing (CIRCLE-Seq) method^3, 5^. We explored the reads of the eccDNA and found that approximately 9,600 reads were normal reads, approximately 6,100 reads were normal breakpoint reads, and approximately 10,000 were breakpoint reads with CTC reads. We found CTC at various lengths (Fig. 5D), indicating the size variation of circular rDNA. Interestingly, we found the longest CTC read, whose length was over 150,000 nt, covered 13 rounds of RDN genes clusters.

Furthermore, we applied CReSIL on a publicly available WGLS datasets of human and mouse cells to screen for eccDNA. CReSIL identified much smaller amounts of the cyclic eccDNA (less than 200 in the individual dataset) from WGLS (Fig. 5E) than from the eccDNA enrichment approach. The high fraction of acyclic regions was possibly derived from other structural variants. Most of the identified eccDNA that had high coverage depth were mostly originated from satellite repeat locations, such as the centromere region, and resulted in variations of the centromere telomere^16^ (Fig. 5F). We found the eccDNA size of approximately 52,000 nt of alpha-like repeats (ALR)/alpha repeats in the class of satellite repeats identified from the human B-Lymphocyte cell NA24385 dataset located on the centromere region of human chromosome 2. Another example found in the mouse dataset of mouse embryonic placenta, ec5, whose size is approximately 38,000 nt (Fig. 5F), contained GSAT_MM repeat in the class of satellite repeats located on the centromere region of mouse chromosome 9. We also found the identified eccDNA harbored whole gene(s) (Fig. 5G). LILRA6, encoding leukocyte immunoglobulin-like receptor subfamily A member 6, and LILRB3, encoding leukocyte immunoglobulin-like receptor subfamily B member 3 were harbored in ec242, with a length of approximately 25,000 nt and coverage depth of 87x, identified in B-Lymphocyte cell NA12878 (Fig. 5G). In another example in the mouse dataset of pancreatic cancer cell lines under a special treatment, we found the whole gene of Mucl2, encoding mucin-like protein 2, harbored in ec38, with a length of approximately 24,500 nt and 207x coverage depth. These examples showed the importance of eccDNA of focal amplification regions that possibly serve a niche of cells.

## DISCUSSION

We presented a bioinformatic software suite package, CReSIL for accurate identification of eccDNA molecules from long-read sequencing data derived from either eccDNA enrichment or WGLS. We have incorporated useful features to assess the function of eccDNA molecules and visualization for deep investigation of the eccDNA architecture. Compared with other bioinformatic tools for identification of eccDNA, CReSIL maintains the highest precision and a performance with lowest F1 score of 0.98 at coverages as low as 3x, which is better than the other tested tools, NanoCircle, eccDNA_RCA_nanopore, ecc_finder, and Flye. NanoCircle has a good precision and well control false positive very well for simple eccDNA, however, NanoCircle has difficulty to control false positive derived from complex eccDNAs. EccDNA_RCA_nanopore was designed to identify eccDNA at the read level and strictly based on the CTC reads; therefore, the tool missed detection of eccDNA that are only represented by non-CTC reads in the sequencing library. Based on the real-world samples (Fig. 3A), the fraction of CTC reads across samples was highly diverse, depending on sample preparation. Therefore, eccDNA_RCA_nanopore missed detection of several eccDNA that had not completed circular amplification by RCA in one amplicon or DNA breakage events. Besides, eccDNA_RCA_nanopore provides the result of individual reads. Therefore, the user needs to write a custom script to reduce the redundancy of the result. Flye in metagenomic mode, which was used in the development of CReSIL software^16^, performed very well when assembling eccDNA at high sequencing depths (F1 score > 0.93 for 10x-100x; the F1 score dropped to 0.73 at the 5x and have difficulty at 3x [Fig. 2F]). This is caused by the design of Flye that is made to assemble genomes; therefore, small sized eccDNA with low sequencing depth are difficult to captured.

Besides high accuracy, CReSIL provides many useful features to study eccDNA. CReSIL generated consensus sequences and variants of individual eccDNA that can be used to assess potential phenotypic effects of eccDNA when variations on the chromosomes are generated. Genes or gene fragments on the eccDNA can be expressed and alter protein levels, resulting in growth advantages for the cells carrying them^24, 38^. Moreover, the eccDNA is also thought to be involved in stable chromosomal gene duplication events when eccDNA reintegrate in the genome^5, 20^. Furthermore, repeats such as alpha-satellite harbored in eccDNA could contribute to centromere variation^16^. Thus, identifying the genetic elements on eccDNA is important to understand of the impact of eccDNA. To aid the user to investigate the eccDNA architecture, CReSIL provides a function to generate the config file, populate the chromosomal origin information and genomic annotations, and plot it in Circos for visualization.

Detection of eccDNA from WGLS challenging^43^; however, it can be possible to uncover the higher copy eccDNA over the genomic background (Fig. 5). As shown, by CReSIL we found that WGLS data be used as basis for identification of eccDNA, when the circular DNA is amplified over the general background level of genomic DNA. This is especially relevant in tumor data where oncogenes are often amplified on large eccDNA^20, 24, 38^. This mean that WGLS of tumors can also be used to identify eccDNA with effects on tumorigenesis, and biobanks of WGLS from tumors serve as a resource for identification of such eccDNA.

CReSIL relies on the reference genome read alignment result, enabling construction of linkages among regions. Therefore, CReSIL cannot capture the eccDNA that originated outside the reference genome sequences that is the limitation of the software. In the future, unmapped reads, especially unmapped CTC reads, can be an additional step to identify eccDNA outside the reference sequences.

In summary, the CReSIL suite software solves major problems in the identification and analysis of eccDNA. It provides a tool for accurate and sensitive identification of eccDNA from long-read sequences, and it thereby allows identification of eccDNA form repeat regions of genomes like the human and for mapping of complex eccDNA made of several fragments from long reads. CReSIL thereby provides a versatile tool to study the immense diversity and impact of eccDNA created from any part of a genome from existing WGS data as well as data enriched for eccDNA.

## METHODS

### Animal and tissue harvest

Wildtype C57BL/6NRj male mice were purchased from Janvier Labs and housed at the University of Copenhagen under ethical approval 2019-15-0201-01659 by Fødevarestyrelsen (The Danish Veterinary and Food Administration) to Jørgen Frank Pind Wojtaszewski. Mice were housed at the University of Copenhagen according to regulations. Three pancreases (12 months old animals IDs 33 and 27 and 3 months old animal ID 27) and three cortexes (3-month old animals IDs 24 and 35 and 12-month old animal ID 38) were used for this experiment. Tissues were stored at -80 °C since the collection date (December 2019). Around 25 mg of tissue were used to extract High Molecular Weight DNA or HMW DNA using MagAttract HMW DNA kit (Qiagen).

### Cells, cultivation, and harvest

The human multiple myeloma cancer cell lines, ARP1, EJM, and JJN3 were provided by Hongwei Xu, PhD and Fenghuang Zhan, MD, PhD (University of Arkansas for Medical Sciences, Little Rock, Arkansas), 5TGM1 murine multiple myeloma cells (RRID:CVCL_VI66) were originally provided by Babatunde Oyajobi, MD, PhD, MBA (University of Texas at San Antonio, San Antonio, Texas), E0771 murine breast cancer cells were obtained from American Type Culture Collection (RRID:CVCL_GR23), and MLOY4 murine osteocyte-like cells (RRID:CVCL_0P24) were originally provided by Lynda Bonewald, PhD (Indiana University, Indianapolis, Indiana). The multiple myeloma cells were cultured in RPMI 1640 culture medium with 10% fetal calf serum, E0771 cells were cultured in DMEM with 10% fetal bovine serum, MLOY4 cells were cultured in alpha-MEM with 2.5% fetal bovine serum and 2.5% bovine calf serum. All cell types were cultured on dishes coated with rat tail collagen type 1 at 37 °C in 5% CO_2_; routinely tested for mycoplasma; and authenticated by morphology, gene expression profile, and for multiple myeloma cells, tumorigenic capacity. Yeast *S. cerevisiae* was cultured in a shake flash in YPD media with 20g/L of glucose. The yeast cells were harvested at the mid-log phase for DNA extraction using NucleoBond HMW DNA (Macherey-Nagel) following the manufacturer protocol.

### EccDNA enrichment

Cells (approximately 10^6^ cells or 10-25 mg of tissue) were washed twice with cold phosphate buffer saline (PBS) pH7.4. Total genomic DNA was then extracted using Monarch Genomic DNA Purification Kit (New England BioLabs) or MagAttract HMW DNA Kit (Qiagen) according to the manufacturer’s protocol. EccDNA was enriched, following the CIRCLE-seq method^12, 38^ with modifications. For each reaction, approximately 5 μg of purified DNA was treated with the genomic rare-cutting endonuclease MssI (Thermo Fisher Scientific) for human samples. For mouse samples either genomic rare-cutting endonuclease SwaI (Thermo Fisher Scientific) or by Cas9 Nuclease, *S. pyogenes* (New England BioLabs) with single guide (sg)RNA1:GTAGCATGAACGGCTAAACGA and sgRNA2: GGCCTGATAATAGTGACGCT to linearized the mitochondrial DNA. The exonuclease V (New England BioLabs) was used to digest linear DNA at 37 °C for 7 days. Every 24 hours, the reaction was filled by adding 30 units of exonuclease V, 10 mM ATP, 13 μL 10x NEBuffer4. During the 5 to 7 days of exonuclease-digested reaction, 2 μL of the exonuclease-digested reaction was tested to confirm the elimination of linear chromosomal and mitochondria DNA by qPCR to amplify chromosomal markers such as the β-globin gene (Hemoglobin subunit Beta)^38^, Cytochrome C Oxidase Subunit 5B^12^, and a mitochondria marker (Cytooxygenase 1)^12^. Digested eccDNA was heat-treated for 20 minutes at 65 °C and purified with 1.8x AMPure XP beads (Beckman Coulter).

### EccDNA amplification and sequencing

RCA was conducted on a heat block at 30 °C for 48 hours using either REPLI-g Mini Kit (Qiagen) or 4BB TruePrime RCA Kit (4basebio). The RCA product was further treated with T7 endonuclease (New England BioLabs) at 37 °C for 1 hour to debranch the product. Debranched DNA was then purified using 1x AMPure XP (Beckman Coulter). DNA concentration was measured using the Qubit dsDNA HS Kit (Thermo Fisher Scientific). Approximately 500 ng of debranched DNA of each sample was used as the input for sequencing library preparation using Native Barcode Kit (Oxford Nanopore Technology [ONT]) following by the manufacturer’s protocol. Then approximately 200 ng of the barcoded DNA (3-5 samples) was mixed and used as input for Ligation Sequencing Kit (ONT) following the manufacturer’s protocol. The summary of the eccDNA enrichment procedure of individual samples is provided in Supplementary Table S3. For yeast whole genome long-read sequencing, the purified DNA of 1 μg was used as the input for Rapid Sequencing Kit (ONT) following the manufacturer’s protocol. All sequencing libraries were performed on a single R9.4 FLO-MIN106D flow cell on a MinION (Mk1B; ONT) for 48 hours through MinKNOW software (ONT) to generate fast5 files.

### Data preprocessing

The fast5 files were basecalled using Guppy software version 5.07 (ONT) with the super high accuracy model (dna_r9.4.1_450bps_sup.cfg) to generate fastq files. The reads with mean a quality score at least 8 and read length at least 200 nt were used for further analysis. We initially aligned reads to the reference genomes and discharged reads that did not align to the reference (see Supplementary Table S4). We investigated the unaligned reads using centrifuge^44^ queried to the non-redundant (nt) database and found the human and mouse reads with low mapping quality (mapQ < 30) and microbes contamination reads (see Supplementary File 1).

### Development of CReSIL software implementation

We recently developed a prototype of CReSIL and NanoCircle^16^ to identify eccDNA from long read sequences that was used as the foundation of this study. CReSIL software was developed and implemented in Python Programming Language version 3. The CReSIL algorithm has 4 major steps plus a generation configuration input for Circos visualization. The input for the software was the fastq files of long-read sequences. It is recommended that CReSIL process the reads with at least 200 nt.

Step 1: Reference-based trimming. CReSIL preprocessed the individual reads to remove “ghost sequences”. Reads were aligned on the reference genome using Minimap2 version 2.24^45^ through mappy.pyx. CReSIL checked for CTCs from the chromosomal alignment locations of individual reads. For the reads containing CTC, CReSIL kept the portion of read that contained the region(s) aligned to the reference genomes with the same alignment orientation (called aligned blocks); the rest of the read was trimmed out. For the reads containing no CTC, CReSIL checked the location and orientation of the aligned block(s). If the read contained more than 1 aligned block, CReSIL kept the first aligned block at the 5’ end of the read (similar to the sequencing direction that begins at 5’ end of the DNA molecule) then extended on the 3’ end to keep the aligned block(s) that had a gap or overlap length of a maximum of 50 nt from the previous aligned block to compensate for micro INDEL events^46^, otherwise the extension stopped and the rest of the read was trimmed out. If the chromosomal alignment location of the aligned blocks overlapped and had the same strand orientation, the aligned blocks were merged, otherwise the longest aligned block was kept. CReSIL generated new trimmed fasta files and chromosomal alignment locations of the trimmed reads that were used for the next steps. (see Supplementary Information: Algorithm 1)

Step 2: Regions and linkages identification. CReSIL merged chromosomal alignment locations of the trimmed reads using bedtools version 2.30 through pybedtools version 0.8.2 to identify chromosomal regions, representing the origin of eccDNA molecules. CReSIL identified linkages among the identified regions from the reads that span over 2 ends of a region or between region(s) known as breakpoint reads (see Supplementary Information: Algorithm 2).

Step 3: Graph construction for eccDNA identification. CReSIL constructed graphs based on the results from Step 2 using networkx version 2.6.3. It was important to ensure that the linkages properly spanned from end to end with the correct orientation. CReSIL represented the individual region with 4 nodes to capture all possible ends and strand information. Then the nodes were connected based on the linkages. The direction of the edges followed the direction of the aligned breakpoint reads. Sequencing a double-stranded DNA can generated 2 reads; therefore, the direction of the edges and the orientation of alignment can be sophisticated, especially the complex eccDNA that needed to be unified in the edge direction. We developed an algorithm to unify the graph by beginning with the longest and walked through the connected path. The unified graphs contained node(s) of represent region(s) and the unified strand(s) and unified edge(s) with weight values, which provided confidence level of linkages. CReSIL checked the individual unified graph for a closed chain of node(s) with circle edge(s) of the same direction to finally identify the eccDNA (see Supplementary Information: Algorithm 3).

Step 4: EccDNA assembly and annotation. We developed an algorithm called region(s) linkage assembly, which is a reference-based assembly approach. CReSIL constructed a draft assembly by extracting the sequence(s) and the orientation of the identified region(s) with respect to the unified graph using Pysam version 0.17.0. For unified graph containing more than 1 node, CReSIL put the extracted sequence in the same order and direction of the unified graph. The draft assembled sequence was then polished by consensus sequencing of the trimmed reads; reads belonging to the region(s) were identified using medeka 1.5.0 and gave eccDNA sequences with variants of SNV and/or INDELS. CReSIL then annotated the genomic features, such as exon, intron, CpG, and repeat, based on a priori genomic annotations. The genomic annotation function of CReSIL allowed the user to use other annotation retrieved from reference genome databases or custom annotation. For demonstration of the CReSIL software, we downloaded an annotation table of the human and mouse genome from UCSC database^47^ consisting of repeats and genes (exon, intron, 5’UTR, and 3’UTR) based on the longest transcript isoform and CpG islands (see Supplementary Information: Algorithm 4).

Step 5: Visualization of eccDNA. CReSIL provided function to populate the results obtained from the CReSIL workflow to generate an input file, including the information of the eccDNA and their genomic annotations (Fig. 1.5), for Circos visualization to present the architecture of individual eccDNA identified by CReSIL.

CReSIL identified eccDNA from WGLS. We extended the capability of CReSIL to identify eccDNA from WGLS by additional workflow. The reads were aligned to the reference genome using minimap2 to generate BAM files. The mosdeth version 0.30 software was used to calculate the depth per window of 10 nt and chromosomal average depth from the BAM file. Next the focal amplified regions were searched throughout the reference genome by selecting only the windows that with a depth greater than the chromosomal average depth. Then, the windows were merged by their chromosomal positions using bedtools2 to produce the focal amplified merge regions. The CReSIL trimming function (Fig. 1.1), was applied to identify the breakpoint reads and their breakpoint locations on the chromosomes. The breakpoint depth that resulted from the pile-up of the breakpoint locations was constructed and used to refine the focal amplified merge regions by trimming the regions outside the breakpoint(s).

The identified regions and their linkages were inputted to the CReSIL graph construction function (Fig. 1.3) and followed the CReSIL standard workflow to identify the eccDNAs.

### Simulated eccDNA datasets for performance evaluation

#### Performance evaluation of eccDNA identification from sequencing data derived from eccDNA enrichment and RCA amplification

We simulated eccDNA sequences by randomly selecting region(s) from the human reference genome with different sizes for 2,600 simple and complex eccDNA sequences to generate a true positive dataset of 1,300 and a true negative dataset of 1,300. To mimic the sequences of RCA and breakpoint(s) for the true positive dataset, we concatenated the sequence of the individual simulated eccDNA five times and used it as a template to generate long-reads with different sequencing depth (3x, 5x, 10x, 20x, 50x, and 100x) using PBSIM2^37^. We manually curated the individual simulated eccDNA to ensure the presence of breakpoint reads in all individual eccDNA. For the true negative dataset, only the simulated eccDNA sequences were used as a template to generate long-reads using PBSIM2 with the same sequencing depth as the true positive dataset. We used F1 scores to evaluate the performance of eccDNA identification of individual tools tailored for eccDNA identification from long-reads such as NanoCircle^16^, eccDNA_RCA_nanopore^15^ (redundant results derived from eccDNA_RCA_nanopore were reduced before the calculation), ecc_finder^29^, and Flye^36^. The results generated by different tools in the fasta file format were converted into a graphical fragment assembly (gfa) format. The fasta files were compared with the simulated eccDNA sequences using dnadiff function of Mummer3^48^. The information of cyclic and acyclic eccDNA was retrieved from the gfa files. The correct detection of eccDNA was defined as cyclic sequences with at least 90% reciprocal overlaps and 90% identity with the simulated eccDNA sequence. The rest was considered as incorrect detection.

#### Performance evaluation of eccDNA identification from whole genome sequencing data

To evaluate the performance of CReSIL in the identification of eccDNA from WGLS data we simulated human chromosomal reads of 20x coverage (∼60 Gb) from the hg19 reference genome using PBSIM2. The simulated human chromosomal reads were mixed with the simulated eccDNA reads (both true positive and true negative datasets) at different eccDNA coverage depth of 5x, 10x, 20x, 50x, and 100x. We used F1 scores to evaluate the performance of eccDNA identification by CReSIL of the individual mixed dataset. The correct detection of eccDNAs was defined as cyclic sequences with at least 90% reciprocal overlaps and 90% identity with the simulated eccDNA sequence. The rest was considered as incorrect detection.

### Publicly available datasets

To demonstrate the capability of CReSIL in the identification of eccDNA from WGLS data, we retrieved the publicly available WGLS datasets of human and mouse genomes from the Sequence Read Archives database. We considered only the WGLS datasets with sequencing amounts greater than 5x of genome coverage. The detail of the datasets is summarized in Supplementary Table S5.

### Software availability

CReSIL 2.0 is freely available at https://github.com/visanuwan/cresil under MIT license.

## Data availability

All data generated in this study will be available at the Sequence Read Archives database under bioproject number PRJNA806866.

## Acknowledgement

We thank Thidathip Wonsurawat for technical discussion and evaluation on DNA debranching. We thank Taylor Wadley for technical assistance on yeast sequencing. We thank Henriette Pilegaard for mice housing, ethical permits, and administration.

## Author Contribution

I.N. design and conceive the project. V.W. and P.J. developed and implement the CReSIL software. V.W., PJ and I.N. performed computational analysis. T.L. and C.M.B performed eccDNA enrichment and sequencing of cell line samples under supervision of I.N. Cell culture experiments were performed under J.D. laboratory. G.A. and M.C.T performed eccDNA enrichment and sequencing of mouse tissues under supervision of B.R. B.R. and G.A developed a protocol for eccDNA enrichment. I.N. wrote the manuscript and all authors edited the manuscript. All authors have read and approved the final version.

